# Integrative profiling of early host chromatin accessibility responses in human neutrophils with sensitive pathogen detection

**DOI:** 10.1101/2020.04.28.066829

**Authors:** Nikhil Ram-Mohan, Simone A. Thair, Ulrike M. Litzenburger, Steven Cogill, Nadya Andini, Xi Yang, Howard Y. Chang, Samuel Yang

**Affiliations:** Department of Emergency Medicine, Stanford University School of Medicine, Stanford, CA 94305; Center for Personal Dynamic Regulomes, Stanford University, Stanford, CA 94305

## Abstract

Sepsis is a leading cause of death globally where neutrophils respond to pathogens via tightly regulated antimicrobial effectors. Combining early neutrophilic responses and pathogen detection may reveal insights for disease recognition. We performed ATAC-seq of human neutrophils challenged with six toll-like receptor ligands and two organisms; and RNA-seq after *Escherichia coli* (EC) exposure for 1 and 4 hours along with ATAC-seq. ATAC-seq of neurophils retains more pathogenic DNA reads than standard library preparation methods. Only a fraction of differential chromatin regions overlap between challenges. Shared signatures exist for ligands but rest are unique in position, function, and challenge. Epigenomic changes are plastic, only ∼500 are shared by EC challenges over time, resulting in varied differential genes and associated processes. We also identify three classes of chromatin mediated gene regulation based on their relative locations. These and transcription factor footprinting reveal timely and challenge specific mechanisms of transcriptional regulation in neutrophils.

## Introduction

Sepsis, a life-threatening sequela of bloodstream infections due to dysregulated host response, is the leading cause of death related to infections worldwide with rising incidences. Most common bloodstream infection causing bacteria are *Staphylococcus aureus* (SA) and *Escherichia coli* (EC) with frequencies of 20.5% and 16% respectively in culture positive patients^1^. Time is of the essence in sepsis, as every hour delay in appropriate antibiotic therapy decreases survival by 7.6%^2^. Understanding early host-pathogen interplay in sepsis can offer invaluable clinical insights critical to saving lives. Neutrophils are the first responders to infection and have been extensively studied for their role in infection and inflammatory processes, particularly sepsis^3^. Neutrophils recognize pathogen associated molecular patterns (PAMPs) via toll like receptors (TLRs)^4^ and danger associated molecular patterns (DAMPs) via receptors such as RAGE for HMGB1^5^. PAMPs are derived from the cell walls of live or dead pathogenic organisms (exogenous signals), whereas DAMPs are derived from the host (endogenous signals) and each are specifically recognized by different TLRs^6^. Both result in inflammatory responses involved in sepsis. Neutrophils responding to PAMPs and DAMPs are capable of unleashing immediate, antimicrobial effector functions, including neutrophil extracellular trap (NET) production, phagocytosis, superoxide production and release of cytokines for further recruitment of other neutrophils and macrophages in a tightly regulated manner^7,8^. Moreover, studies have described that the neutrophil life span may be extended from 5-8 hours in the periphery to days upon interaction with both PAMPs and DAMPs^9,10^. These responses are tightly regulated to avoid collateral damage like increased vascular permeability and hypotensive shock resulting from release of heparin binding proteins by neutrophil activation via adherence to endothelial cells^11^ and lung injury and poor patient outcome because of a cytokine storm resulting from hyperresponsiveness and dysregulation of apoptosis in lung neutrophils^12^.

Despite possessing tightly regulated yet diverse functions, neutrophils have been regarded as a terminally differentiated cell type with limited ability to produce transcripts or proteins. The inference from this assumption is that the chromatin structure of a neutrophil is dynamically limited. Specifically in comparison to monocytes, neutrophils were shown to have much lower gene expression and largely repressed chromatin^13^. However, despite the fact that neutrophils have reduced transcriptional activity overall, they possess a much more dynamic range of transcripts and CpG patterns when compared to other cell types^14^. Neutrophils also show heterogeneity in their methylation patterns between individuals^15^ and undergo active chromatin remodeling, methylation/acetylation patterns associated with gene transcription and cytokine production^16–19^. Neutrophils employ an inhibitor program to safeguard their epigenome from unregulated activation thereby protecting the host^20^. Epigenetic signatures have also been shown to play a role in the cellular function of septic patients^21^. Specifically, bacteria can affect the chromatin structure of host immune cells via histone modifications, DNA methylation, restructuring of CTCF loops, and non-coding RNA^19,22,23^. Such chromatin changes allow for repositioning of inflammatory genes into a transcriptionally active state, recruitment of cohesion near the enhancer regions, and result in swift transcriptional response in the presence of EC^19^. Even though chromatin remodeling is shown in neutrophils in response to external stimuli, the exact regulation of transcription by changes in chromatin accessibility is not well understood.

Since the epigenome reacts before gene expression, we are interested in profiling chromatin responses in neutrophils to infections for early disease recognition. In the current study, we f i r s t explore the relevant chromatin elements involved in TLR-mediated responses to 1hour exposures from various pathogen ligands or whole SA and EC on a genome-wide scale using ATAC-seq to elucidate differences in the host response. We also assay the temporal fluctuation between 1 and 4 hours post EC exposure in the neutrophil epigenome and resulting transcriptome to better understand the processes and pathways involved in immune response. Neutrophilic chromatin accessibility patterns may reveal what pathogen an individual has encountered and/or how they are responding to the infection. Our analyses reveal chromatin accessibility, enriched motifs, and functional signatures unique to each challenge. We also observe time specific chromatin accessibility changes resulting in transcriptional changes at two time points. Based on the chromatin accessibility patterns, we classify three categories of transcriptional regulation in neutrophils. Additionally, since prokaryotes lack chromatin, employing ATAC-seq results in increased pathogen to host ratio of DNA, enhancing rare microbial reads. The coupling of host response profiles with microbial reads in a single assay may offer diagnostic advantages while gaining unprecedented insights into early host-pathogen dynamics and neutrophil biology.

## Results

### Neutrophils are activated in response to ligand and whole organism challenges

Purified neutrophils from 4 female healthy volunteers were challenged with an array of toll-like receptor (TLR) agonists for one hour, namely lipotechoic acid (LTA) (TLR2), lipopolysaccharide (LPS) (TLR4), flagellin (FLAG) (TLR5), resiquimod (R848) (TLR7/8); as well as β-glucan peptide (BGP) that signals via the dectin-1 receptor^24^ for fungal infection and the DAMP high mobility group box 1 (HMGB1), a cytokine released in sterile inflammation (such as early traumatic events, thought to signal through RAGE and TLR4)^5,25^ (Fig. 1b). Neutrophil activation was confirmed by IL8 and TNFα qRT-PCR (Fig. 1c). Since it is important that the ATAC-seq is performed on intact nuclei, sytox green assay was performed to estimate the extracellular DNA as an indication of NETs. No NETs were observed in response to any stimuli at the time of ATAC-seq (Fig. 1d).

**Fig. 1.**
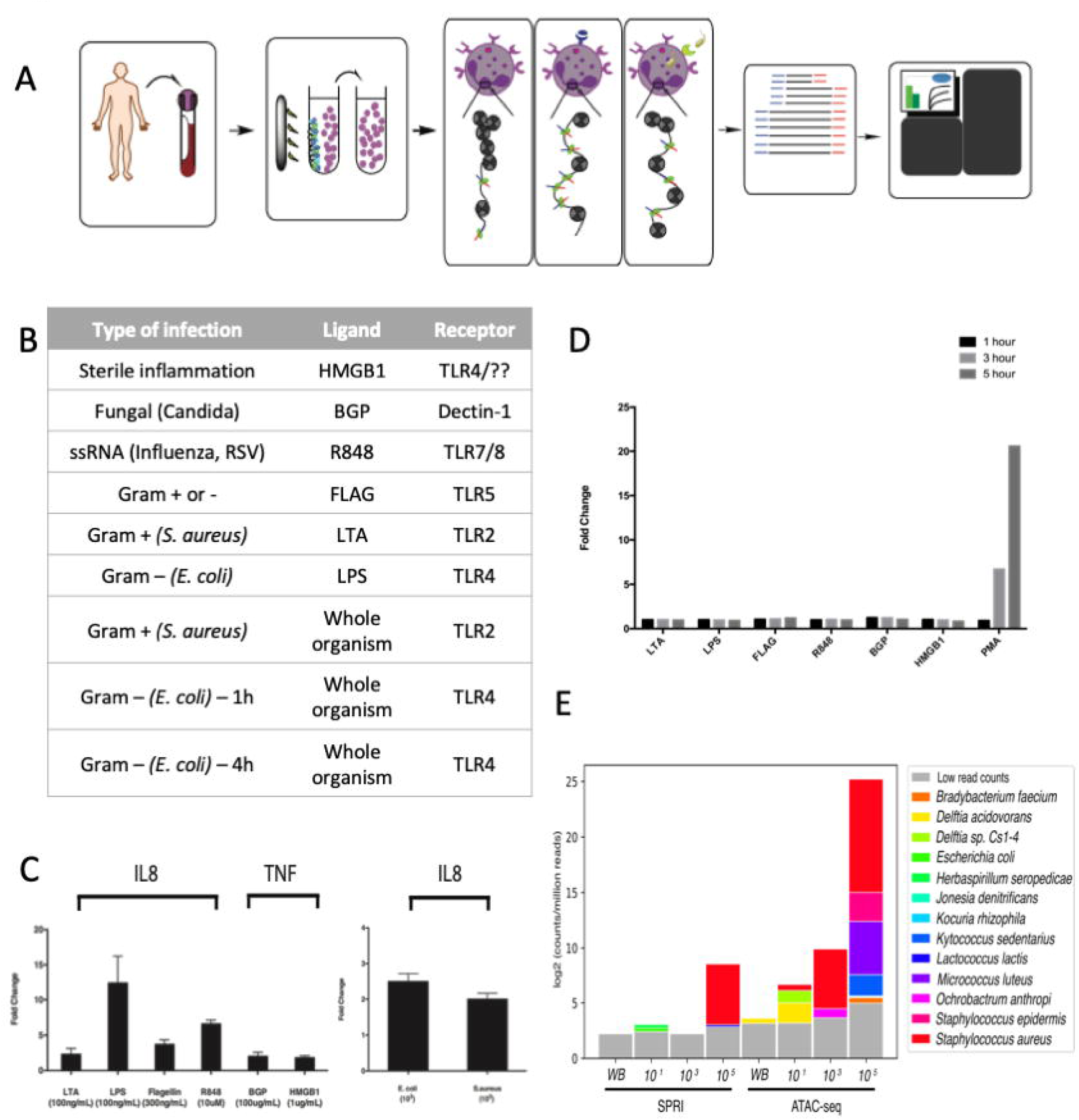
Neutrophil activation in response to challenges. **a.** Schematic of neutrophil isolation, stimulation, and ATAC-seq. Blood is collected from healthy volunteers in EDTA tubes and unwanted cells are removed using a magnetic bead selection. Tn5 transposase (green ovals) carrying an adaptor payload (red and blue) complementary to next generation flow cells and inserts randomly into regions of open chromatin. Unstimulated and stimulated neutrophils are the sequenced using Illumina technology. **b.** Table of tested challenges including 6 ligands, 2 whole organisms, and 1 time series. **c.** Healthy donor neutrophils produce IL8 or TNF in response to ligands or live organism challenge (ligand donors n= 4, live organism donors n= 2, mean and SE are represented. **d.** Healthy volunteer neutrophils do not produce NETs via sytox green assay in response to pathogen ligands at 1 hour or immediately following live organism challenge supporting this time point for ATAC-seq. (PMA is a positive control) (ligand donors n=4, live organism donors n=2). **e.** ATAC-seq is more sensitive to SA reads than traditional SPRI library preparation. Whole blood was spiked at increasing concentrations with live SA and prepared for sequencing using traditional DNA extraction and library preparation (SPRI method) compared to neutrophil isolation and ATAC-seq method. (n = 2) (WB: whole blood only, no organisms). Relative abundance plots illustrate that reads align to SA (red) as well as other bacteria, however the species that would be contamination are still low in relative abundance.

### Pathogen DNA from challenges is enriched in ATAC-seq

Neutrophils possess largely closed chromatin, and prokaryotes lack chromatin. ATAC-seq on neutrophils yields several features that increase sensitivity for pathogen reads. First, negative isolation of neutrophils reduces the number of human cells and potentially captures any circulating or phagocytized pathogens. Second, by surveying only open chromatin, ATAC-seq increases the pathogen to host ratio of DNA in the sample compared to traditional library preparation methods. To demonstrate this, neutrophils were challenged with SA in incremental colony forming units (CFU) per mL for 1 hour. These were then parallelly subjected to genome-wide sequencing using a standard SPRI library preparation and ATAC-seq. As suspected, the relative abundance of SA reads obtained by ATAC-seq was higher than the SPRI method, for all concentrations (Fig. 1e). In fact, the relative abundance retained by ATAC-seq at 10^3^ CFU/mL is comparable to the 10^5^ CFU/mL SPRI preparation method; a marked improvement in sensitivity. Contaminant signals are present given that the neutrophils were only challenged with SA, these are likely short, low complexity reads that that do not map specifically. Despite the contamination, ATAC-seq samples display 3 times more reads for the pathogen compared to SPRI.

### Differential accessibility of chromatin in the genome is challenge and time specific

Neutrophils have limited accessible chromatin; however, insert size distributions and enrichment at transcription start sites (TSS) were consistent across samples (Supplemental Figs. 2a and 2b). Strong correlation of genome wide peak counts across technical replicates (from r^2^ = 0.70 - 0.95) were observed for any given ligand stimulation (Supplementary Fig. 3) while lower correlation was observed between donors, suggesting donor specific heterogeneity.

Differential accessibility of chromatin in neutrophils is readily apparent when comparing the accessible chromatin landscape in response to the challenges with that of unstimulated neutrophils using Diffbind. For each challenge, filtering of the differential region calls with p < 0.05 and abs(logFC)>=1 resulted in a total of 28,812 differential regions for the LTA challenge; 35,453 for with LPS; 34,038 with FLAG; 33,604 for R848; 29, 198 for BGP; 32,916 with HMGB1; 10,201 with SA; 31,197 with the EC*-*1h (EC1h) exposure; and 22,511 with EC-4h (EC4h) exposure. Despite the differences in the numbers of differential regions in each challenge, their genomic distribution identified using ChIPseeker (Fig. 2a) is similar. More than 80% of the differential regions in each challenge were found in either distal intergenic or intronic regions.

**Fig. 2.**
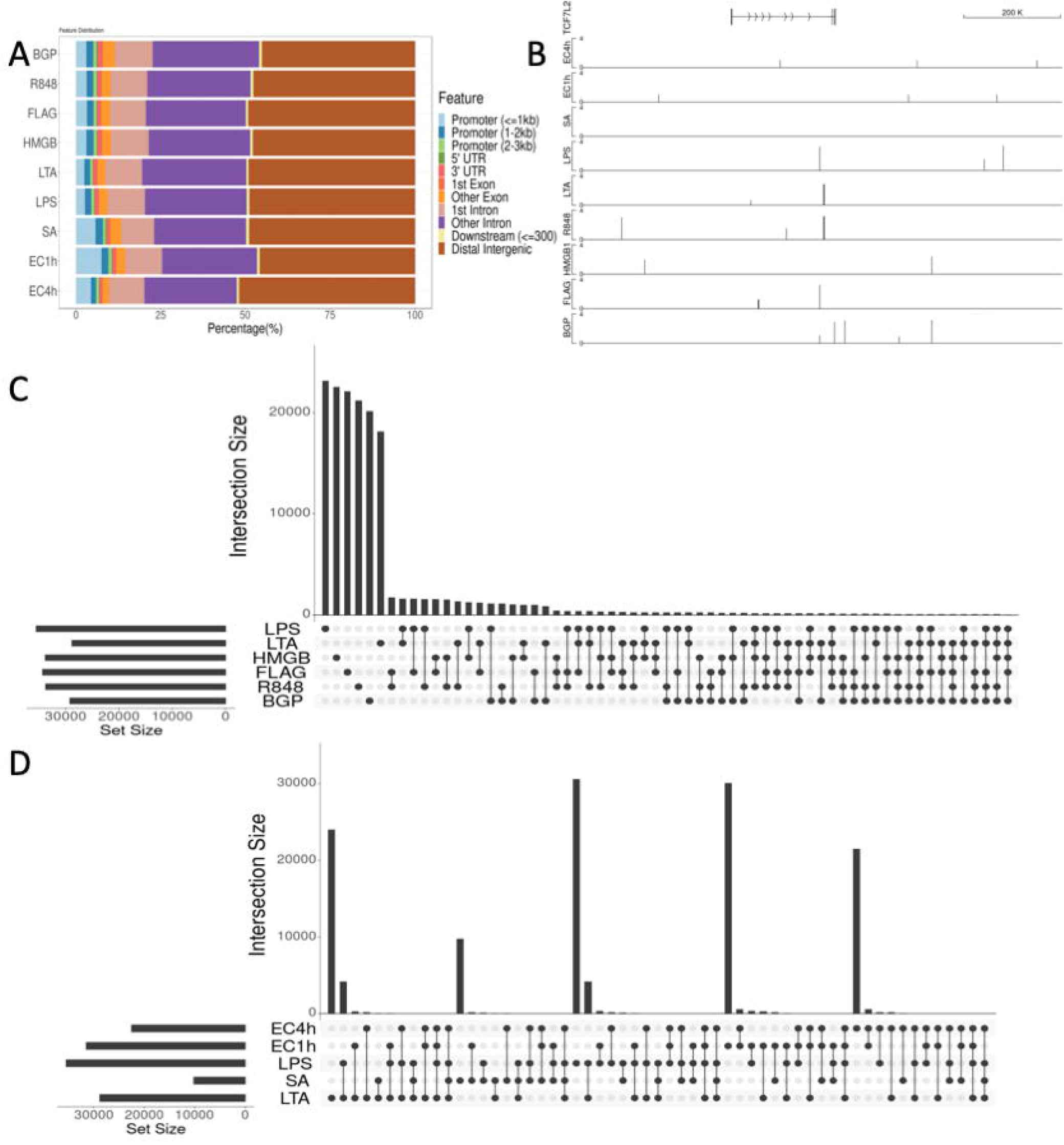
Comparison of locations of differential regions across challenges. **a.** Whole genome distribution of differential regions within promoters, UTRs, exons, introns, downstream regions, and distal regions as determined using ChIPseeker. Very similar distribution patterns across sample, more than 80% of differential regions are distal. **b.** Depiction of the unique signatures across the challenges. Differential regions around the TCF7L2 gene. From top, all transcripts generated by the TCF7L2 gene, the TCF7L2 gene, differential regions associated at EC4h, EC1h, SA, LPS, LTA, R848, HMGB1, FLAG, and BGP respectively. **c.** Upset plot of overlapping differential regions across ligand challenges. Consensus set of regions were generated using Diffbind and the presence absence was visualized using Upset on R. Majority of the differential regions are unique to challenges. **d.** Upset plot of overlapping regions between the whole organism challenges and their corresponding ligands. Minimal overlap between whole organisms and ligands.

Landscape of differential chromatin accessibility is challenge dependent. This uniqueness is down to the gene level where associated differential regions between challenges are disparate (Fig. 2b). A map of the differential regions around the TCF7L2 gene portrays this uniqueness clearly. Overlap analyses of differential regions across the entire genome between challenges showed that the vast majority of these regions did not overlap and were challenge specific (Figs. 2c and 2d). A total of 19,710; 21,597; 21,978; 22,712; 17,791; 20,703 regions were unique to the tested ligands BGP; FLAG; HMGB1; LPS; LTA; and R848 respectively. Similar patterns were observed with the whole organism challenges as well. In the SA; EC1h; and EC4h challenges, there were 9,417; 28,818; and 20,766 unique regions respectively. There are no overlapping differential regions across all of the challenges. 6 out of the 9 challenges tested were the most to have any overlapping differential regions. A total of 115 regions were shared by a combination of 6 challenges, 603 were common between 5 challenges, 2121 between 4 challenges, 6429 between 3 challenges, and 23,149 regions were common between a combination of two challenges. Interestingly, out of the 115 differential regions common to 6 challenges, 108 are common across all the ligand challenges (Fig. 2c). There is only 1 common differential region between the whole organism challenges. However, comparing the signature between the whole organism challenge and their corresponding ligands, a few commonalities exist (Fig. 2d). There are 72 common differential regions between the SA and LTA challenges, 548 common between the two EC time points, EC1h and LPS have 356 in common, and EC4h and LPS have 188 in common. Surprisingly, however, only 6 differential regions are common to the two EC time points and LPS. On average, ∼64% of the differential regions from the merged peaksets are unique to the ligand challenges while with the whole organism challenges, ∼92.3% of the regions are unique to the challenge.

Unique chromatin accessibility signatures in response to challenges are not limited to specific positions of differential chromatin accessibility but correspond to differences in the regulated functional pathways as well. Differential regions were assigned to genes by using a combination of prediction tools and surveying overlaps with known regulatory regions. This was done rather than assigning regions to genes immediately downstream so as to account for distal regulation as well. Prediction methods included T-gene^26^ and GREAT^27^. Overlap analyses were performed against the HACER database as well as the predicted regulatory regions identified in primary human cancers^28^. Applying this combinatorial method, a minimum, ∼91.6% of the differential regions were associated with genes while on average ∼95.6% of the differential regions were successfully associated with genes across all challenges. In addition, on average, 78.3% of these overlapped previously identified regions with histone marks from the IHEC. GOterm and Pathway enrichment analyses with the assigned genes and the number of differential regions associated with each of them revealed dissimilar functional enrichment signatures between challenges (Fig. 3a). Signaling by receptor tyrosine kinases is the only common pathway enriched amongst all 9 challenges. Metabolism of carbohydrates, NOTCH1 signaling pathways, and deactivation of beta-catenin transactivating complex are unique to the whole organism challenges. The EC challenge over time resulted in similar pathway enrichments of the differentially accessible chromatin regions. Pathways like TLR4 cascade, neutrophil degranulation, response to infectious diseases, and Fc gamma receptor dependent phagocytosis are enriched. Overall, other than for a few overlapping enriched pathways, each challenge elicited varied functional responses upon stimulating the neutrophils.

**Fig. 3.**
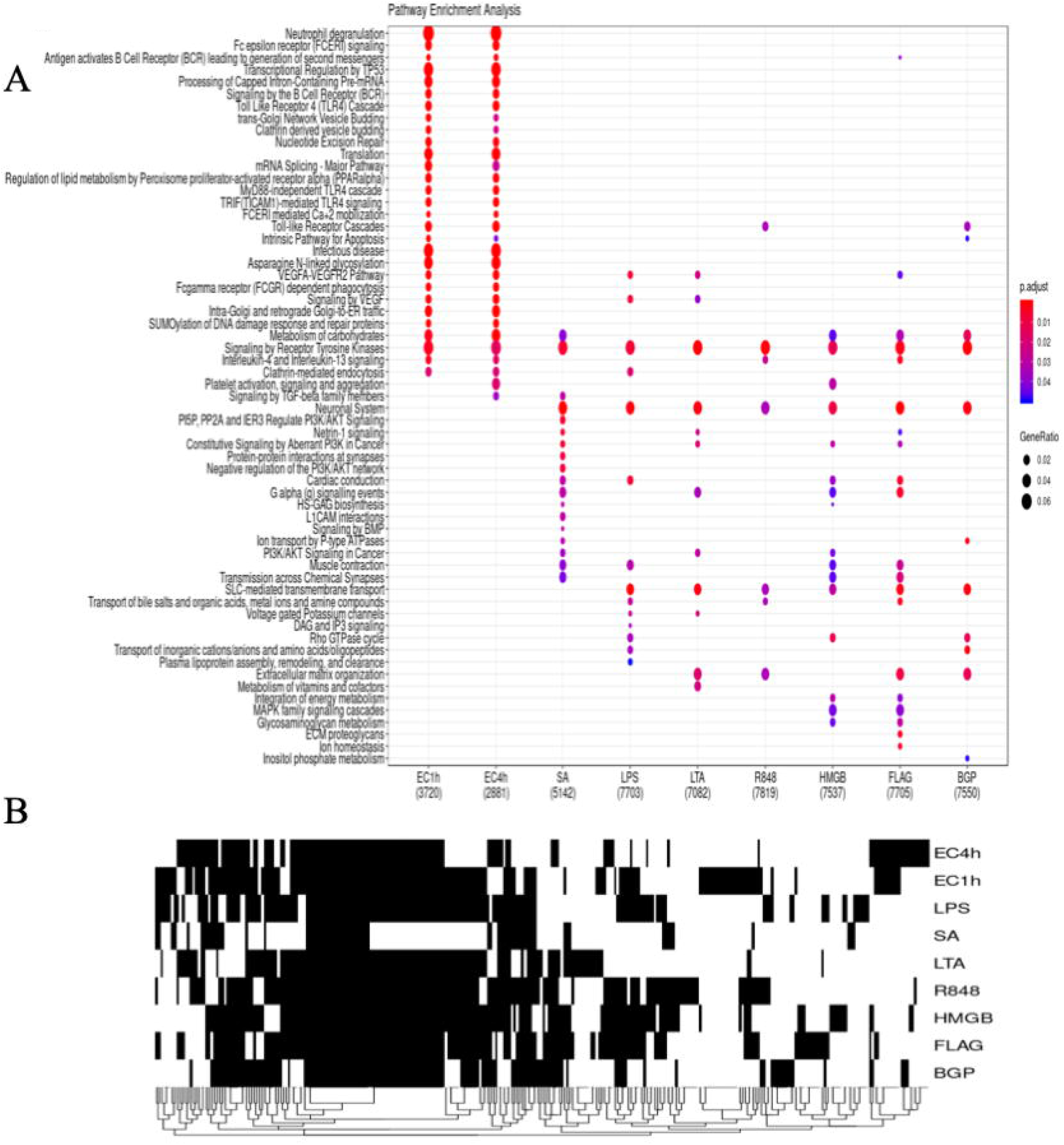
Comparison of functional profiles of differential regions. **a.** Reactome pathway enrichment analysis of the genes associated with the differential regions in each challenge. Gene assignments to each peak were carried out as described in the methods. Enrichment analysis performed using ChIPseeker and compared using the comparecluster function from clusterprofiler. **b.** Presence-absence heatmap of enriched motifs in the differential regions from each challenge. Enriched motifs were determined using HOMER and filtering for a p-value of p<0.01. Although a lot of enriched motifs are shared, there are motifs unique to each challenge.

Similar to patterns with position and function, there are unique enriched motifs specific to each challenge. Importantly however, despite large proportions of differential regions being unique to each challenge, these still contain common enriched motifs shared across challenges. For example, even though there are no common differential regions across all 9 challenges, 24 enriched motifs are common to them (Fig. 3b). Additionally, comparing enriched motifs in all the differential regions across the challenges, there are motifs unique to each. There were 5 unique motifs in the BGP challenge; 6 in the FLAG challenge; 5 in HMGB1; 5 in LPS; 6 in LTA; 7 in R848; 2 in SA; 14 in EC1h; and 5 in EC4h (Supplemental table 1). Hence, it is important to combine both specific positions as well as the enriched motifs to identify signatures unique to each challenge.

### Transcriptional plasticity of neutrophils in response to EC challenges

RNA-seq analysis of EC challenged neutrophils at two time points – 1 hour (EC1h) and 4 hours (EC4h) using edgeR revealed a temporal pattern in gene expression suggesting plasticity in neutrophil transcription. Although most genes remain unchanged in expression in the presence of EC, there are differences in the number of differentially expressed genes and the magnitude of change between the two time points (Fig. 4a). There are 66 up and 56 down regulated genes at 1 hour of exposure while there are 2554 up and 2656 down regulated genes at 4 hours. There are only 93 genes up regulated and 10 genes down regulated common to both time points. Interestingly, there are a few genes that are either up regulated at 1 hour and down regulated at 4 hours or vice versa. In this category, 7 genes were up at 1 hour and down at 4 and 10 were down at 1 hour and up regulated at 4 hours.

**Fig. 4.**
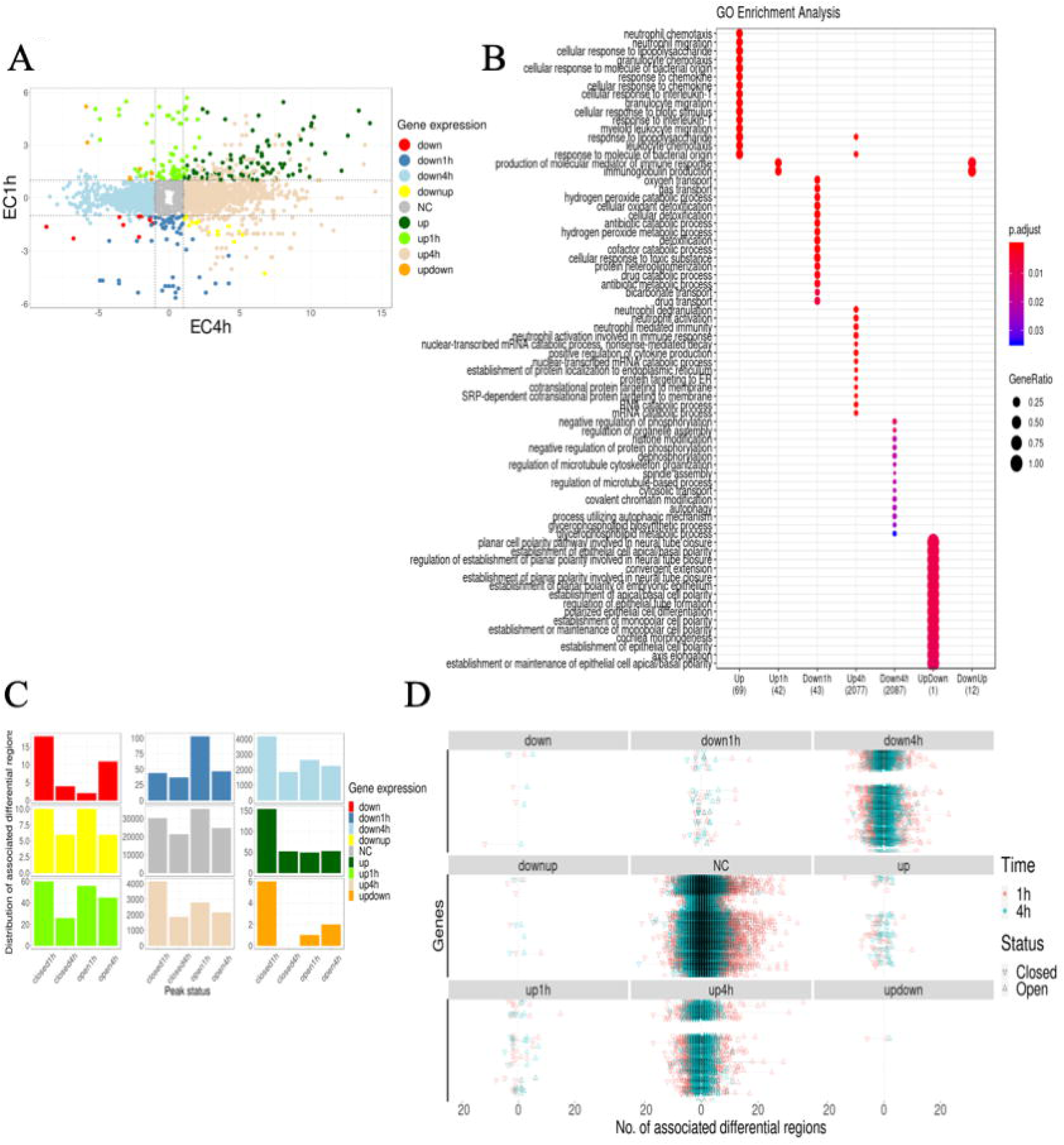
Paired ATAC-seq and RNA-seq of neutrophils challenged with EC at 1 and 4 hours. **a.** Comparison of expression of genes at the two time points. logFC calculated using edgeR for each gene at the two time points were plotted as points if the p value is less than 0.05. Points are colored based on the gene expression patterns at the two time points. down: genes down-regulated at both time points; down1h and down4h: genes down-regulated only at 1 hour and 4 hour time points respectively; downup: genes down-regulated at 1 hour and up regulated at 4 hours; NC: genes that are not differentially expressed at either time point; up: genes up-regulated at both time points; up1h and up4h: genes up-regulated only at 1 hour and 4 hour time points respectively; and updown: genes that are up-regulated at 1 hour and down-regulated at 4 hours. **b.** GOterm enrichment analysis of the genes categorized based on their pattern over time. Grouped genes analyzed using the enrichGO feature within clusterprofiler. **c.** Distribution of open and closed differential regions at each time point with respect to the gene expression patterns. Combination of open and closed regions at each time point and each gene expression pattern except for genes that are up regulated at 1 hour and down regulated at 4 hours. **d.** Counts of open and closed differential regions associated with each gene at each time point. Typically, more associated differential regions for each gene at 1 hour than at 4 hours. Additionally, combination of multiple open and closed differential regions associated with most genes.

Plasticity is also reflected in the biological processes enriched at the two time points (Fig. 4b). GOterm enrichment of genes grouped in above mentioned categories portrays a transcriptional landscape in neutrophils that are responsive to external stimuli. For example, immunoglobulin production is transiently up regulated at 1 hour, suggesting immediate response to the challenge. Transport and metabolism are predominant in the down regulated genes at 1 hour. Similarly, at 4 hours of EC challenge, processes involved in neutrophil activation and degranulation, response to molecules of bacterial origin, intrinsic apoptotic signaling pathway, and processes utilizing autophagy mechanisms are enriched in the up regulated genes. More diverse processes are enriched in the down regulated genes at 4 hours and interestingly, many overlaps with those enriched in the up regulated genes at 4 hours. As expected, however, many processes involved in immune response are enriched in the genes that are up regulated at both time points. These include neutrophil migration, cellular response to lipopolysaccharides, positive regulation of inflammatory response, chemokine mediated signaling, NF-kappaB signaling, and regulation of DNA binding transcription factor activity. Surprisingly there are no enriched processes in the continuously down regulated genes. Interestingly, SRP-dependent co-translational protein targeting to membrane was down regulated at 1 hour and up regulated at 4 hours. On the contrary, Fc-epsilon and Fc-gamma receptor signaling pathways, regulation of complement activation, and regulation of protein processing are all up regulated at 1 hour and down regulated at 4 hours.

### Transcriptional plasticity of neutrophils is a result of complex accessible chromatin crosstalk

Combination of ATAC-seq and RNA-seq revealed a complex combination of differentially open and closed chromatin regions affecting gene expression changes. For each category of gene expression, a count of the whether the associated peaks are differentially closed or open at each time shows that there is no specific pattern (Fig. 4c). That is, genes being up regulated at 1 hour are not all a result of just differential chromatin regions either opening or closing. For genes that are differentially expressed only at 1 hour, there likely are chromatin accessibility changes occurring at 4 hours that are maintaining gene expression level with the control. This is readily apparent when looking at the distribution of the openness or closedness of associated differential regions (Fig. 4c). This phenomenon is clear when surveying the associated differential regions for each gene (Fig. 4d). For example, out of the 7 genes that are up regulated at 1 hour and down regulated at 4 hours, differential chromatin regions were found to be associated only with 1 gene, PTK7. For this gene, 9 associated differential chromatin regions either open or close in a combinatorial manner to affect the gene expression. These 9 associated regions include 6 regions that are differentially closed at 1 hour, 1 that is differentially open at 1 hour and 2 that are open at 4 hours. Interestingly however, there is no direct correlation between gene expression and the number of associated differential regions (Spearman correlation – EC1h: 0.0321 EC4h: 0.0289). In general, there are more chromatin accessibility changes earlier while transcriptional changes largely occur at the later time point.

Based on the *in silico* linked chromatin accessibility changes and the transcriptional expression, the regulatory mechanisms were classified into three proposed categories – 1. Differential regions only in the promoter (1h: 84; 4h: 86); 2. Differential regions only in distal sites while the promoter region is primed for expression (1h: 19,450; 4h: 23,206); and 3. Differential regions in the promoter as well as distal sites (1h: 2245; 4h: 1015). Although these three mechanisms are prevalent, there are differentially expressed genes that do not have any associated differential chromatin regions. In the first category, Differential regions were present only in the TSS ± 2.5 kb regions (Fig. 5a). An example for this category is the FAM66C gene which is affiliated with the long non-coding RNA (lncRNA) class. This de novo differential opening in the promoter region results in ∼3 logFC increase at 4 hours in the presence of EC. In the second category, regions in the promoter are open, however, there is no difference between the EC challenge in comparison to control. In this category, gene expression is fine-tuned by distal regulatory regions. The BBS2 gene, a member of the Bardet-Biedl syndrome gene family and forms a part of the BBSome multiprotein complex, is down regulated at 4 hours (−1.4 logFC) while there exists an associated differentially open region present at 1 hour. This region being open facilitates the maintenance of gene expression at 1 hour (Fig. 5b). The third category is a combination of differential chromatin regions in the promoter region and in associated distal sites. An example is the HMGCS1 gene that encodes a protein with protein homodimerization, and isomerase activity is up regulated ∼3 logFC only at 4 hours (Fig. 6a). Interestingly, the accessible chromatin signature associated is complex. A differential chromatin region is fixed in the promoter at both time points and yet differential gene expression occurs only at one. This is likely a result of the associated distal regions – two that are differentially open at 1 hour and one each of differentially open and closed regions at 4 hours. These distal interactions are further supported by the Hi-C predicted interacting regions^**19**^. These distal regions fall within 226,569 Hi-C predicted interactions across the genome. A complex interplay of interacting chromatin regions facilitates expression of the associated gene. These interactions facilitate the activation or repression of binding motifs to fine tuning the regulation. Transcription factor footprinting of enriched motifs in open distal differential regions associated with the HMGCS1 gene revealed a temporal pattern of binding (Fig. 6B). The Sp2 binding motif remains unbound at both time points while the ETS1 motif is only bound at 1 hour and the n-Myc motif is bound only at 4 hours. Similar patterns of binding are observed in motifs unique to each time point at these differential regions as well. A combination of varied differentially accessible chromatin regions and transcription factor binding motifs provide an intricate means of transcriptional regulation in neutrophils.

**Fig. 5.**
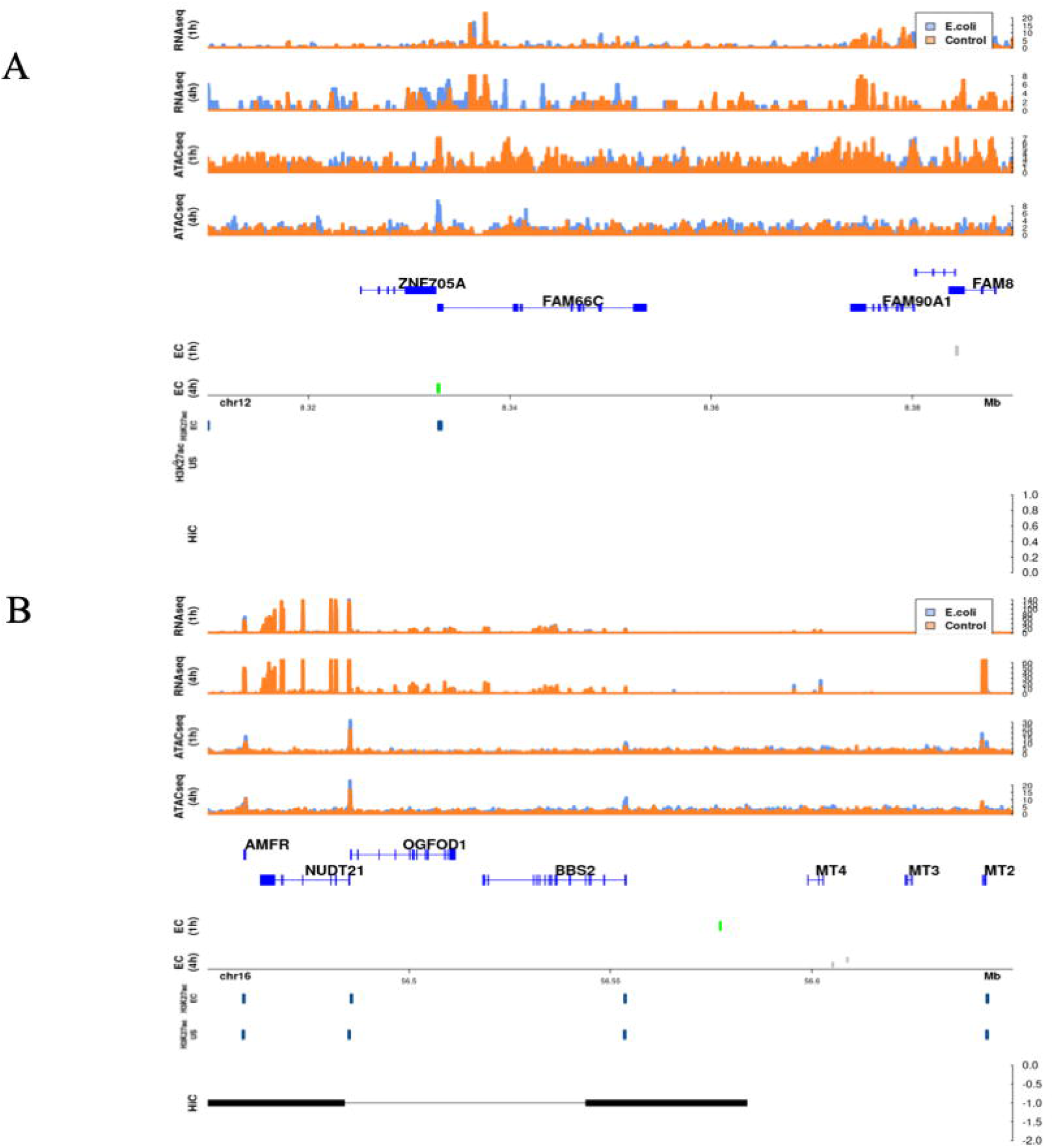
Mechanisms of control of transcription by accessible chromatin in neutrophils. **a.** Differential regions only in the promoter. Promoter is defined as the regions ±2.5Kb around the TSS for each gene. Example for this category is the FAM66C gene. **b.** Differential regions only in distal regions, promoter is primed for expression. Example for this category is the BBS2 gene. In A and B, from the top, RNA-seq coverage at 1 hour for EC and control; RNA-seq coverage at 4 hours; ATAC-seq coverage at 1 hour; ATAC-seq coverage at 4 hours; genes in the region from hg19; open (green) differential regions associated with the gene at 1 hour and open or closed differential regions not associated with the gene of interest (gray); differential regions associated at 4 hours; location on the chromosome; H3K27ac histone marks in the presence of EC; and absence of EC lifted over from earlier study^19^; and Hi-C interactions in the presence of EC lifted over from earlier study^19^.

**Fig. 6.**
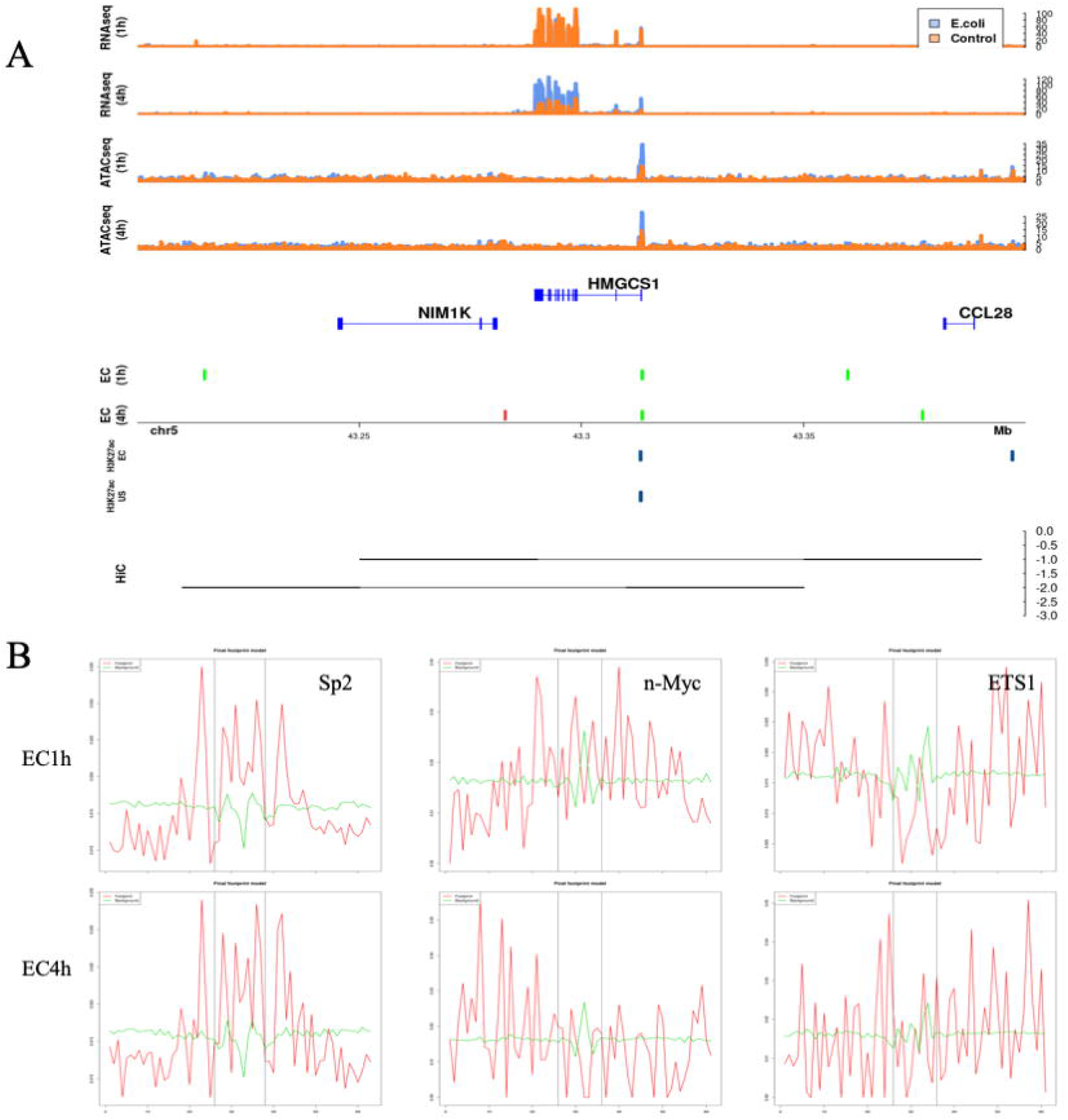
Intricate combinatorial chromatin accessibility regulation of transcription in neutrophils. **a.** Third mechanism of gene regulation in neutrophils involves differential regions in both the promoter as well as distal sites. An example in this category is the HMGCS1 gene. The differential region in the promoter is fixed and unique to an EC infection. Tracks are similar to those in Fig. 5. Differentially closed regions are shown in red **b.** Transcription factor footprinting of enriched motifs identified in the distal associated differential regions. Footprinting using the ATAC-seq reads was performed using the FootprintingPipeline. Time dependent binding of transcription factors affecting gene expression.

## Discussion

In this study we explore the chromatin accessibility signatures affecting immune response in neutrophils challenged with external infecting stimuli using ATAC-seq. ATAC-seq on neutrophils provides a unique advantage, interrogating the host response biology, while simultaneously gathering information about pathogens with higher detection sensitivity. Identifying the pathogen responsible for sepsis is an area of intense research. Additionally, ATAC-seq can evaluate the accessibility changes in intergenic and enhancer regions^29^, showing cell type specific enhancer usages. Even though next generation sequencing has provided technological advances, gathering useful host and microbial information simultaneously is challenging. Only a fraction of the reads generated f rom human samples correspond to pathogens and, even if successfully retained in the sea of human DNA, contig assembly is very difficult, so classification must be able to proceed accurately with short reads from noisy data^30,31^. There is a large body of work dedicated to defining positivity via setting thresholds (adjusted based on pathogen load and misclassification due to incomplete reads), species identification to differentiate pathogen from contaminant, and subtraction from negative controls^30,31^. Our assay design improves sensitivity by capitalizing on negative isolation of neutrophils, reducing the amount of background human DNA as well as cell free microbial DNA in the final sequencing sample and then only sequencing open chromatin from the eukaryote and prokaryote. It performs drastically better than standardized diagnostic library preparation methods (Fig. 1e). This assay requires a small sample volume making it ideal towards clinical adoption in sepsis diagnosis.

Neutrophils are known to form subpopulations at the site of inflammation in response to the stimulus^32^ but it was unclear whether epigenetic changes are challenge specific. We find challenge specific genome wide chromatin changes (Figs. 2b, 2c, and 2d) despite similar genomic distributions (Fig. 2a). These unique chromatin changes reflect in common and unique enriched pathways as well specific transcription factor binding motifs unique to each challenge (Fig. 3). Only signaling by receptor tyrosine kinases is common to all challenges. This supports the challenge specific response nature since the only common pathway is sensing external cues. Although there are only about ∼500 differential regions common to the two EC challenge time points, their associated enriched pathways are identical (Fig. 3a). Interestingly however, there aren’t many shared pathways between EC and the corresponding ligand, LPS portraying the differences between single ligand and whole organism stimulation. Apart from the receptor tyrosine kinase pathway, VEGFA signaling pathways is shared which is involved in neutrophil recruitment^33^. These chromatin accessibility signatures support the earlier discovered stimulus specific gene expression changes in response to LPS and EC^34^. The unique chromatin accessibility signatures also expose unique transcription factor binding motifs specific to challenges (Fig. 3b and supplemental table 1).

Early events in immune cells involved in sepsis can and should be captured in their epigenome, as this is the first step in the cellular response to a cell’s environment. These chromatin accessibility events could be a source for new diagnostic tools and even novel molecular targets for new therapies. At one hour, under many stimuli in this first responder cell type, we find challenge specific genome wide changes in chromatin accessibility (Figs. 2b, 2c, and 2d). This phenomenon is also readily apparent when comparing the differential chromatin accessibility at the two time points of the challenges (31,197 at 1h vs 22,511 at 4h). Measurements of gene and protein expression capture events much later than epigenomic changes and hence may be less informative. For example, it has been shown that enhancer profiling was better at determining cell identity than mRNA^35^. This delayed transcription vs epigenetics is supported by our data. While more chromatin accessibility changes are observed at 1h, more differential expression of genes occurred at 4h and in a time specific manner (Figs. 4b, 4c and 4d). Hence, we propose that a combination of unique differentially accessible chromatin regions as well as motif signatures we have identified may be more illuminating of the neutrophil’s pathogen exposure. These exposure-specific chromatin accessibility changes are rapidly induced and, while many maybe transient, may leave a longer-lasting “mark” on the epigenome, potentially spawning a new forensic and diagnostic modality, an advantage over current tools, as the epigenome is the earliest detectable signal.

Plasticity in neutrophils has been widely accepted recently^36,37^ but the mechanisms driving this have yet to be successfully delineated. Current focus on the role of epigenetics has vastly expanded the understanding^38^, but much is still unknown. A study of unchallenged neutrophils from healthy volunteers identified over 2000 genes with a significant epigenetic component explaining their expression^39^ and the role of epigenetics in sepsis induced immunosuppression in various immune cells has been identified^36^. Subsequently, significant chromatin restructuring was observed in response to a three hour EC infection^19^. While these chromatin restructures facilitate the opening of the inflammatory response armament, the exact accessible chromatin interactions are still unknown. With the EC infection time series in our study, we find that both transcription and chromatin accessibility are plastic in neutrophils. This is readily apparent in the low overlap in the differential regions between the two time points. This can be attributed to the changes in chromatin structure and the need for opening/closing of transcription factor binding sites giving rise to transient gene expression. Based on the differential regions and gene associations, we classified three broad categories of accessible chromatin regulation that occur within earlier identified CTCF anchored loops^19^. The categories include – differential regions only in the promoter, differential regions only in the distal enhancer regions, and finally genes regulated by a combination of both. Of these categories, genes with the differential regions only within the promoter were the fewest (Fig. 5a) and showed new H3K27ac modifications within the promoter. Genes with primed promoter regions being regulated only by distal enhancer regions were the highest (Fig. 5b) and showed histone marks in the promoter, suggesting promoter activation, under both challenged and unstimulated states further supporting our classification. The third category is a combination of both differential regions in promoters as well as distal regions and the HMGCS1 gene, implicated in the regulation of inflammation^40^, is an example of how this results in gene expression (Fig. 6a). Although similar mechanisms of epigenetic transcriptional regulation are known in macrophages^41,42^ and the additive or competitive roles of multiple distal enhancers for gene are known^43,44^, this is the first evidence for such regulation in neutrophils. We also see time specific binding of transcription factors (Fig. 6B) within the HMGCS1 associated distal regions, possibly involved in cooperative activation or repression similar to other systems^45–48^ or by inhibiting the binding of different transcription factors^20^ to result in the observed gene expression. Overall, we show that neutrophils undergo plastic transcriptional expression under intricate accessible chromatin regulation that is unique to the stimulus faced (Fig. 7) and that this methodology can potentially be used in combination with these signatures as a putative diagnostic tool.

**Fig. 7.**
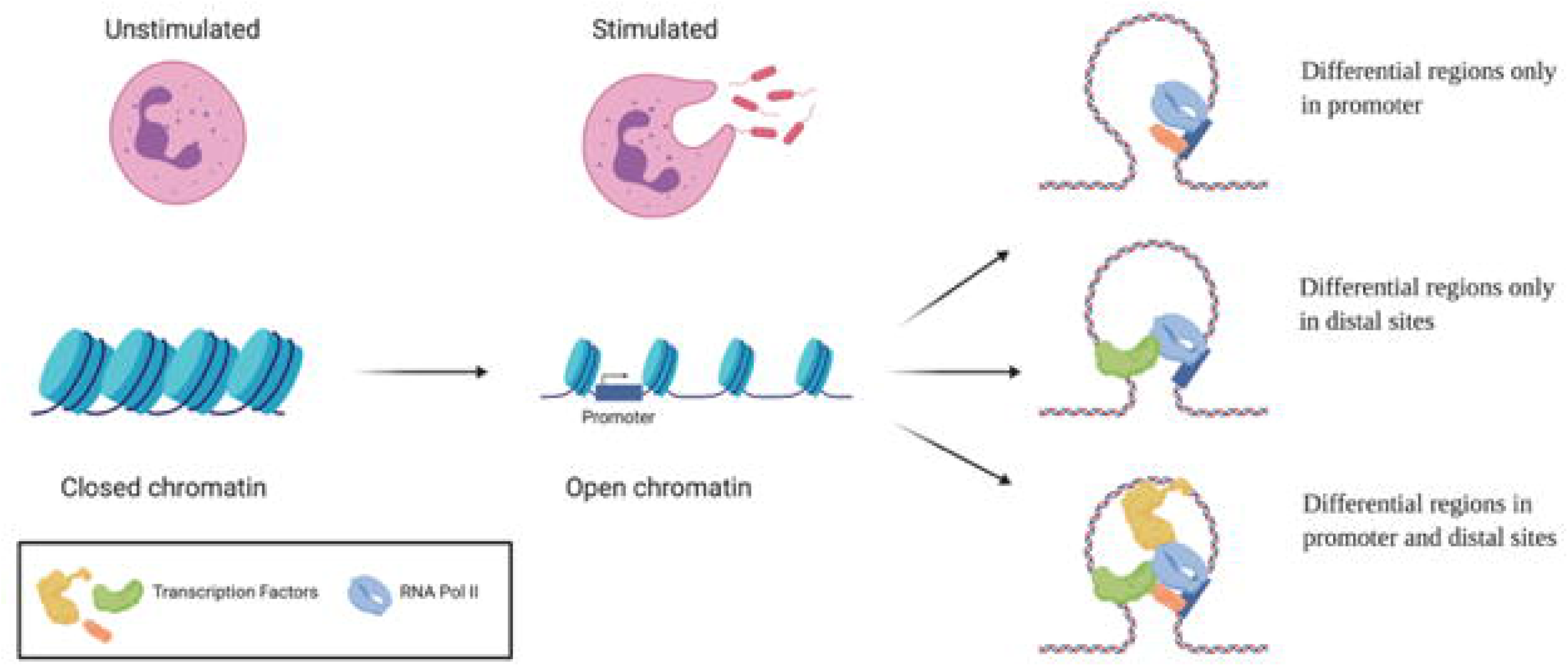
Model for intricate accessible chromatin regulation of gene expression in neutrophils. A schematic depicting the three mechanisms of regulation identified in our analyses. Upon exposure to stimuli, the closed chromatin in neutrophils opens up in stimulus specific patterns. Accessible chromatin regulation of genes occurs in one of three mechanisms where – 1. Differential regions occur only in the promoter region; 2. Differential regions occur only at distal cites; and 3. Differential regions occur at promoter and distal sites.

## Methods

### Study participants

Four healthy volunteer females 30-40 years old were recruited and informed consent obtained (Stanford University IRB-37618).

### Negative selection isolation and activation of neutrophils

#### FACs and Flow cytometric analysis

Neutrophils were isolated using a negative selection method that allowed for seamless dovetailing with the ATAC-seq method (Fig. 1a, Supplemental Fig. 1). Neutrophils were isolated using the Stem Cell EasySep Direct Human Neutrophil Isolation Kit as per the manufacturer’s protocol and resuspended in PBS with 1% FBS and 2mM EDTA. 8.0 × 10^5^ cells per condition were fixed in 1% PFA for 10 min at room temperature and subsequently stained with primary antibodies: mouse anti-human PEcy7-CD16 (BD #557744), mouse anti-human PerCPcy5.5-CD66b (Biolegend # 305107), and mouse anti-human V500-CD45 (BD # 560779) as well as the corresponding IgG controls. Flow cytometric and statistical analysis were performed using FlowJo V. 10.0.8.

### Toll Like Receptor (TLR) samples

Neutrophils isolated from 4 healthy volunteers were plated at 50,000 cells per well and stimulated in duplicate with the following ligands for 1 hour: lipotechoic acid (LTA) 100ng/mL (Invivogen), LPS (Sigma) 100ng/mL, Flagellin (FLAG) 300ng/mL, resiquimod (R848) (Invivogen), 10uM, CpG Class C ODN 2395 5uM,, B-glucan peptide (BGP) 100ug/mL (Invivogen), high mobility group box 1 (HMGB1) (R & D) 1ug/mL^4,5,24,25^ (Fig. 1b).

### Live Organism Challenge samples

Blood from 2 healthy volunteers was spiked with a specific CFU/mL of either EC ATCC 25922 or SA ATCC 29312. The Stem Cell EasySep Direct Human Neutrophil Isolation Kit was applied as per the manufacturer’s protocol to 2mL of blood after 1 hour of SA treatment and 1 and 4 hours of EC treatment. After isolation, cells were counted, divided into 50,000 cell samples (50uL) in duplicate (Fig. 1b).

### Quantitative RT-PCR of IL8 and TNFα

Total RNA was isolated from 8.0 × 10^5^ cells prepared as above with the RNeasy kit (QIAgen). RNA was DNase treated using the TURBO DNA-free DNase treatment (Ambion). One step qRT-PCR was performed in the Rotor Gene Q using the Rotor Gene SYBR Green RT-PCR kit. ΔΔCt was calculated using GAPDH. Primer sets are as follows: *IL8 F 5’CAGTTTTGCCAAGGAGTGCT, IL8 R 5’ACTTCTCCACAACCCTCTGC, TNF F 5’GCTGCACTTTGGAGTGATCG, TNF R 5’ATGAGGTACAGGCCCTCTGA, GAPDH F 5’TGCACCACCAACTGCTTAGC, GAPDH R 5’GGCATGGACTGTGGTCATGA*

### Sytox Assays for Neutrophil Extracellular Traps

Neutrophils were isolated as above using either the TLR sample or live organism sample preparation as appropriate. Cells were plated at 2.0 × 10^5^ per well in triplicate. A positive control was created by stimulating cells with 25nM PMA (Sigma). 5mM Sytox green (Life Technologies) was used to detect the presence of NETs^49^. Fluorescence intensity was measured using the Tecan Infinite M200 Pro.

### ATAC-seq and RNA-seq library prep and sequencing

All treated and untreated control cells from 4 donors and 6 ligands as well as 2 donors and 2 whole organism challenges were collected as described above and ATAC-seq was performed as described^29^. Excess primers libraries were removed using the AMPpure bead kit. In parallel with ATAC-seq, genome-wide sequencing using a standard SPRI^50^ library preparation was performed on the SA challenged neutrophils with AMPure XP from Beckman Coulter.

RNA was extracted from isolated neutrophils after the 1 and 4 hour EC challenges as well untreated controls using the miRNeasy Micro kit from Qiagen and libraries were generated using KAPA PolyA enrichment mRNA library prep. All libraries were sequenced at the Stanford Functional Genomics Facility on the Illumina HiSeq.

### ATAC-seq analysis

#### Data processing and peak calling

The datasets generated for this study are available under BioProject - xxxxxxxx. Fastq files were analyzed from raw data all the way to peak calls using the PEPATAC pipeline (http://code.databio.org/PEPATAC/) against the hg19 build of the human genome. Briefly, reads were trimmed of adapters using Trimmomatic^51^ and aligned to hg19 using Bowtie2^52^ with the –very-sensitive -X 2000 parameters. Duplicates were removed using PICARD tools (http://broadinstitute.github.io/picard/). Reads with MAPQ <10 were filtered out using Samtools^53^. Reads mapping to the mitochondria or chromosome Y were removed and not considered. Technical replicates were merged using Samtools yielding one sample per donor per stimulation. Peaks were called using MACS2^54^ with the -q 0.3 --shift 0 –nomodel parameters. Given the closed nature of neutrophil chromatin and our interest only in differential regions, we chose to relax the FDR cutoff to produce sufficient regions for further differential studies. Correlation between replicates was generated and a single peakset was generated across replicates for each challenge.

#### Microbial identification

Reads generated from both SPRI as well as ATAC-seq were preprocessed by trimming with Trimmomatic. Using Kraken^55^, human reads were removed from the samples, and relative abundances for pathogens were determined as counts per million reads. Replicates were averaged together and log2 transformed for an abundance value.

#### Differential analysis

Differentially accessible regions were identified from the merged peaksets using the DiffBind R package^56^. A p-value of 0.05 and abs(logFC) > = 1 were set as the threshold. Consensus bed files were generated with Diffbind with a threshold of 0.66 overlap. Overlapping/ common regions between peak sets were determined using the DiffBind tool and visualized with the UpSet package^57^ in R.

#### Assigning genes associated with differential regions

Differential regions in each sample were associated with genes following multiple approaches – 1. T-gene^26^ from the MEME suite was used to predict regulatory links between the differential region and the genes. Only associations with p-value < 0.05 and correlation >= 0.4 were included. 2. Using GREAT^27^ and implementing the basal plus extension algorithm and defining a 2.5Kb region each for proximal upstream and downstream respectively, and a distal region up to 500Kb. 3. Surveying overlap of differential regions with previously reported association links^28,58^ using Bedtools^59^. Additional support for these associations was derived by incorporating predicted Hi-C interactions from EC stimulation of neutrophils for 3 hours^19^. A functional pathway enrichment analysis was performed using the ChIPseeker^60^ package on R with the custom developed table with the differential regions and their associated genes. This uses a hypergeometric model to assess the enrichment of genes associated with a pathway. A Benjamin-Hochberg adjusted p-value of 0.01 was used as a cutoff.

#### Annotating differential regions

Genomic distribution of differential regions and enrichment around transcription start sites were estimated using ChIPseeker. A custom background of histone marks was collected from the International Human Epigenome Consortium (http://ihec-epigenomes.org). We selected for mature neutrophil samples and women of Northern European ancestry. Sample files in bigbed format were converted to bed format using the UCSCtools package. Motif analysis was performed using HOMER^61^ and known motifs with a cutoff of p < 0.01 were selected. Each differential region was annotated with known overlapping histone marks and a list of motifs. Presence-absence heatmaps of the enriched motifs in each sample were plotted using heatmap.2 from within the gplots package in R. Footprinting of transcription factors was enriched in regions of interest was performed using the FootprintPipeline (https://github.com/aslihankarabacak/FootprintPipeline).

### RNA-seq analysis

#### Data processing and differential expression analysis

Quality of the paired-end reads generated for each replicate was performed using FastQC^62^ and trimmed with Trim Galore (https://github.com/FelixKrueger/TrimGalore). Resulting reads were aligned to hg19 using HISAT2^63^ with the --rna-strandness RF parameter. The generated SAM files were sorted and then converted to BAM using Samtools. Counts were generated using the R package Rsubread^64^ in a strand specific manner. Differential gene expression analysis was performed using edgeR^65^ and genes with FDR corrected p-value < 0.05 and logFC >= 1 or logFC <= -1 were selected. GOterm enrichment analysis was performed and comparisons between time points were made using the compareCluster function from the cluterProfiler R package^66^ which uses a hypergeometric model to assess the enrichment of genes associated with a pathway. A Benjamin-Hochberg adjusted p-value of 0.01 was used as a cutoff.

## Data Availability

The datasets generated for this study are available under BioProject - xxxxxxxx.

## Acknowledgements

We would like to thank the Stanford Functional Genomics Facility for performing the sequencing carried out in this study. All bioinformatics analyses were performed on the SCG cluster of the Stanford Research Computing Center. Would also like to thank Shin Lin M.D., Ph.D, Assistant Professor of Cardiology at UW for critical review and feedback of the manuscript. Supported by RM1-HG007735 (to H.Y.C.). H.Y.C. in an Investigator of the Howard Hughes Medical Institute and a Postdoctoral Research Fellowship from Stanford Child Health Research Institute to S.T.

## Author Contributions

S.T., S.Y., and H.Y.C. designed the project. S.T. performed the ATAC-seq and N.A. and X.Y. performed the RNA-seq. N.R., S.T., U.L. and S.C. performed all bioinformatics analysis and generated the figures with critical feedback from S.Y. and H.Y.C. N.R. wrote the manuscript with feedback/input from all other authors.

## Figure legends

**Supplemental Fig. 1.** Flow cytometry with gating strategy depicted confirms 98.1% purity of CD66b/CD16 double positive neutrophils.

**Supplemental Fig. 2. a.** Representative QC plots demonstrating library prep results in expected insert size distribution and **b.** reads are enriched around transcription start sites (TSS).

**Supplemental Fig. 3.** Quality control for DiffBind method of identification of differentially accessible regions of chromatin. Correlation heat maps and principal component analysis (PCA) of differentially accessible chromatin. We found that for any given challenge across donors, stimulated samples cluster together, control samples cluster together, and the stimulated and control cluster away from each other, suggesting high quality data and accessible chromatin region identification that allows for analysis of four healthy donor data.

**Supplemental Table 1.** List of known motifs identified using Homer that are unique to each challenge.

