## Supplemental material for "Integrative profiling of early host chromatin accessibility responses in human neutrophils with sensitive pathogen detection"

### Supplemental Figure 1

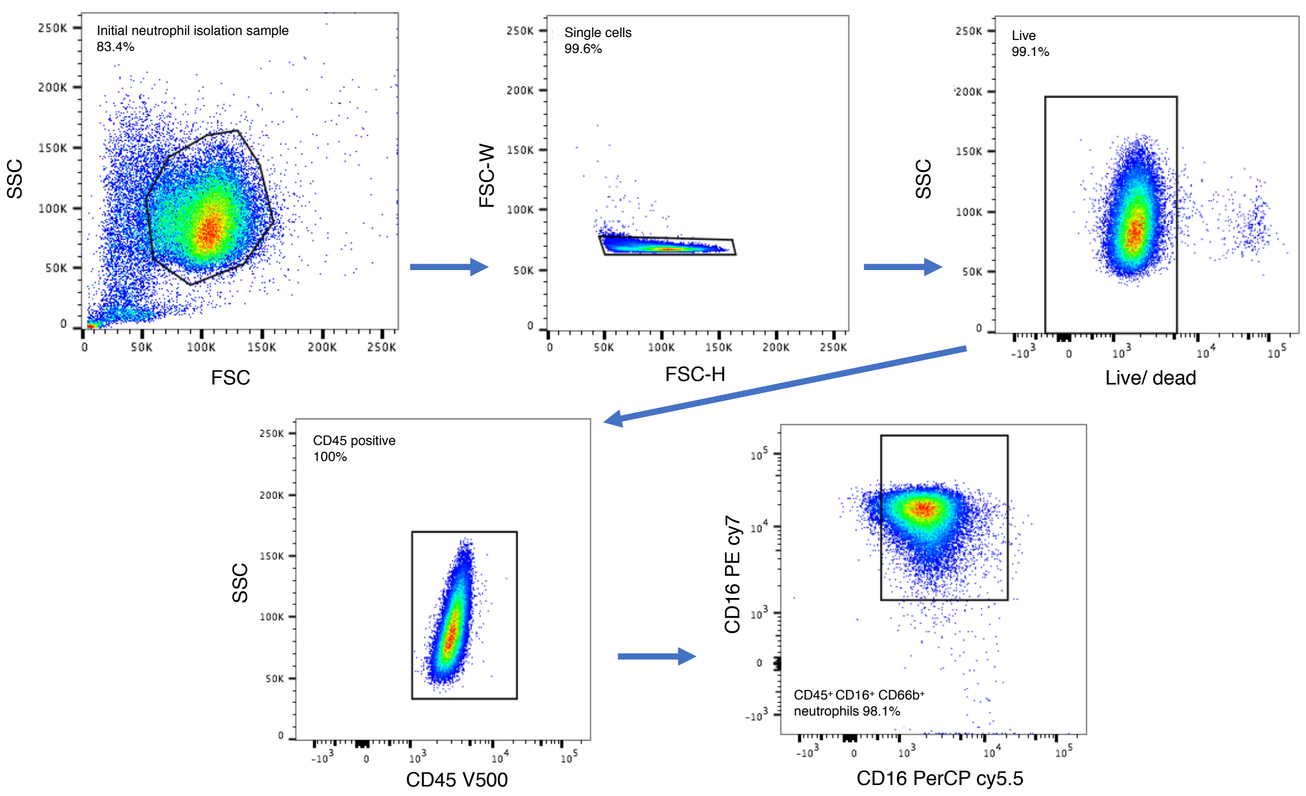


**Supplemental Figure 1.** Flow cytometry with gating strategy depicted confirms 98.1% purity of CD66b/CD16 double positive neutrophils.

**Supplemental Figure 2**

A.

B.

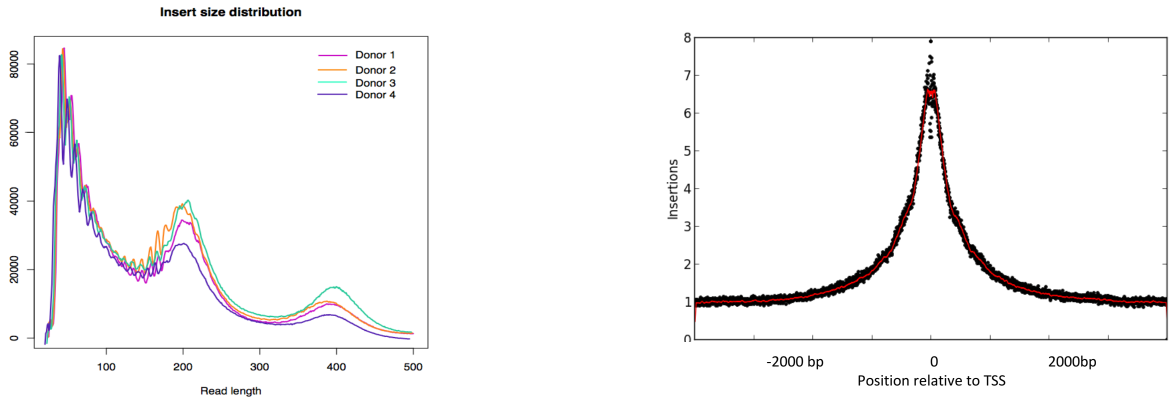


**Supplemental Figure 2**. **A.** Representative QC plots demonstrating library prep results in expected insert size distribution and **B.** reads are enriched around transcription start sites (TSS).

**Supplemental Figure 3.**

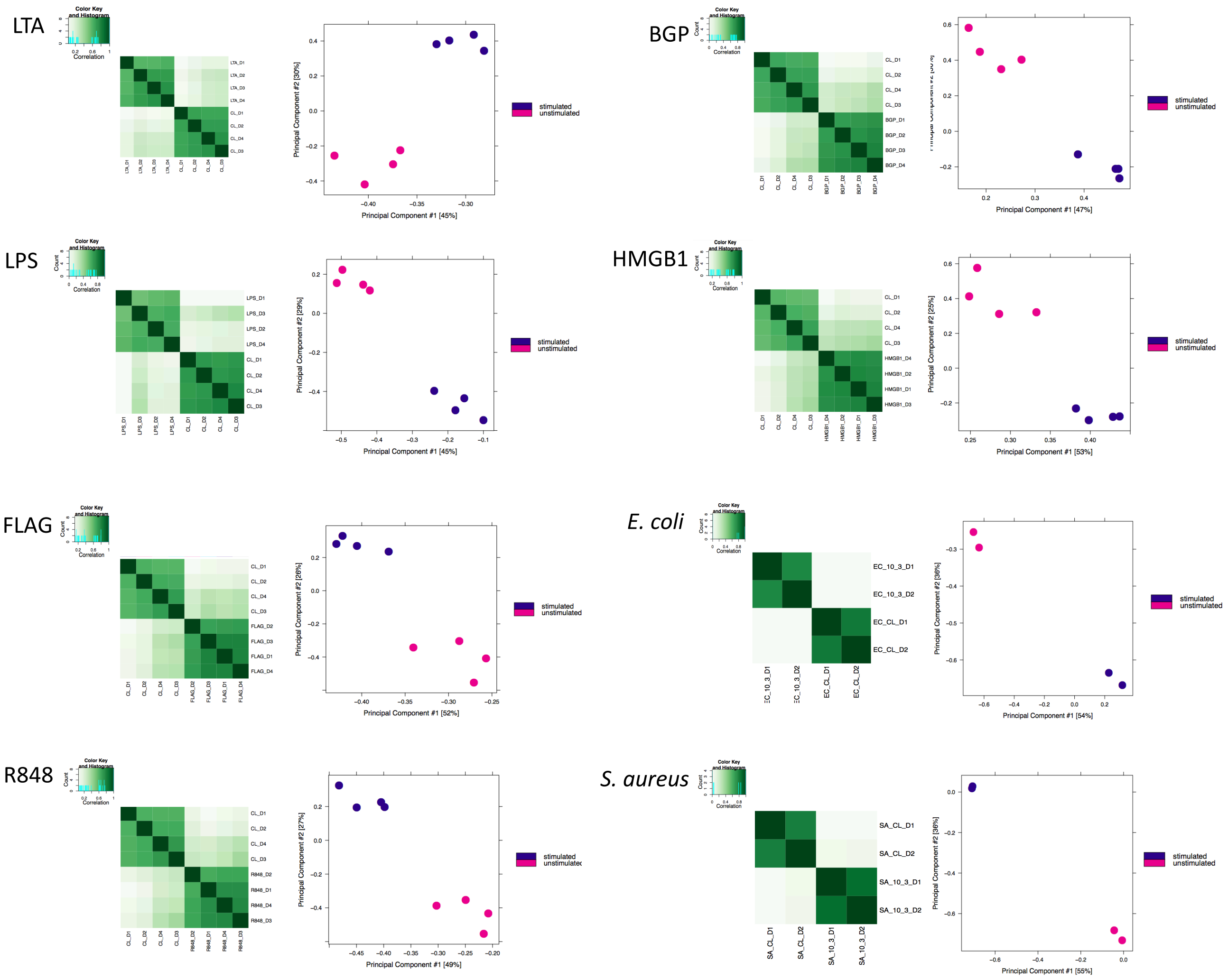


**Supplemental Figure 3.** Quality control for DiffBind method of identification of differentially accessible regions of chromatin. Correlation heat maps and principal component analysis (PCA) of differentially accessible chromatin. We found that for any given challenge across donors, stimulated samples cluster together, control samples cluster together, and the stimulated and control cluster away from each other, suggesting high quality data and accessible chromatin region identification that allows for analysis of four healthy donor data.

Supplemental Table 1.

| Homer-motif | Challenge |
|---|---|
| CEBP:CEBP(bZIP)/MEF-Chop-ChIP-Seq(GSE35681)/Homer | BGP |
| p53(p53)/Saos-p53-ChIP-Seq(GSE15780)/Homer | BGP |
| p53(p53)/Saos-p53-ChIP-Seq/Homer | BGP |
| Rfx2(HTH)/LoVo-RFX2-ChIP-Seq(GSE49402)/Homer | BGP |
| Six2(Homeobox)/NephronProgenitor-Six2-ChIP-Seq(GSE39837)/Homer | BGP |
| T1ISRE(IRF)/ThioMac-Ifnb-Expression/Homer | BGP |
| Egr1(Zf)/K562-Egr1-ChIP-Seq(GSE32465)/Homer | EC1h |
| Egr2(Zf)/Thymocytes-Egr2-ChIP-Seq(GSE34254)/Homer | EC1h |
| GFY(?)/Promoter/Homer | EC1h |
| GFY-Staf(?,Zf)/Promoter/Homer | EC1h |
| GRHL2(CP2)/HBE-GRHL2-ChIP-Seq(GSE46194)/Homer | EC1h |
| HRE(HSF)/Striatum-HSF1-ChIP-Seq(GSE38000)/Homer | EC1h |
| ISRE(IRF)/ThioMac-LPS-Expression(GSE23622)/Homer | EC1h |
| KLF3(Zf)/MEF-Klf3-ChIP-Seq(GSE44748)/Homer | EC1h |
| KLF5(Zf)/LoVo-KLF5-ChIP-Seq(GSE49402)/Homer | EC1h |
| Klf9(Zf)/GBM-Klf9-ChIP-Seq(GSE62211)/Homer | EC1h |
| Ronin(THAP)/ES-Thap11-ChIP-Seq(GSE51522)/Homer | EC1h |
| STAT1(Stat)/HelaS3-STAT1-ChIP-Seq(GSE12782)/Homer | EC1h |
| STAT5(Stat)/mCD4+-Stat5-ChIP-Seq(GSE12346)/Homer | EC1h |
| ZBTB12(Zf)/HEK293-ZBTB12.GFP-ChIP-Seq(GSE58341)/Homer | EC1h |
| Cdx2(Homeobox)/mES-Cdx2-ChIP-Seq(GSE14586)/Homer | EC4h |
| CDX4(Homeobox)/ZebrafishEmbryos-Cdx4.Myc-ChIP-Seq(GSE48254)/Homer | EC4h |
| KLF10(Zf)/HEK293-KLF10.GFP-ChIP-Seq(GSE58341)/Homer | EC4h |
| NFIL3(bZIP)/HepG2-NFIL3-ChIP-Seq(Encode)/Homer | EC4h |
| Prop1(Homeobox)/GHFT1-PROP1.biotin-ChIP-Seq(GSE77302)/Homer | EC4h |
| bHLHE41(bHLH)/proB-Bhlhe41-ChIP-Seq(GSE93764)/Homer | FLAG |
| E2F(E2F)/Hela-CellCycle-Expression/Homer | FLAG |
| Foxf1(Forkhead)/Lung-Foxf1-ChIP-Seq(GSE77951)/Homer | FLAG |
| GATA(Zf),IR4/iTreg-Gata3-ChIP-Seq(GSE20898)/Homer | FLAG |
| Meis1(Homeobox)/MastCells-Meis1-ChIP-Seq(GSE48085)/Homer | FLAG |
| NFAT:AP1(RHD,bZIP)/Jurkat-NFATC1-ChIP-Seq(Jolma_et_al.)/Homer | FLAG |
| HRE(HSF)/HepG2-HSF1-ChIP-Seq(GSE31477)/Homer | HMGB |
| Pax7(Paired,Homeobox),long/Myoblast-Pax7-ChIP-Seq(GSE25064)/Homer | HMGB |
| Sox15(HMG)/CPA-Sox15-ChIP-Seq(GSE62909)/Homer | HMGB |
| Stat3+il21(Stat)/CD4-Stat3-ChIP-Seq(GSE19198)/Homer | HMGB |

| | |
|---|---|
| Tcf7(HMG)/GM12878-TCF7-ChIP-Seq(Encode)/Homer | HMGB |
| CUX1(Homeobox)/K562-CUX1-ChIP-Seq(GSE92882)/Homer | LPS |
| GATA3(Zf),DR8/iTreg-Gata3-ChIP-Seq(GSE20898)/Homer | LPS |
| TR4(NR),DR1/Hela-TR4-ChIP-Seq(GSE24685)/Homer | LPS |
| VDR(NR),DR3/GM10855-VDR+vitD-ChIP-Seq(GSE22484)/Homer | LPS |
| ZNF669(Zf)/HEK293-ZNF669.GFP-ChIP-Seq(GSE58341)/Homer | LPS |
| ETS:E-box(ETS,bHLH)/HPC7-Scl-ChIP-Seq(GSE22178)/Homer | LTA |
| HIF2a(bHLH)/785_O-HIF2a-ChIP-Seq(GSE34871)/Homer | LTA |
| Six1(Homeobox)/Myoblast-Six1-ChIP-Chip(GSE20150)/Homer | LTA |
| STAT6(Stat)/Macrophage-Stat6-ChIP-Seq(GSE38377)/Homer | LTA |
| ZNF16(Zf)/HEK293-ZNF16.GFP-ChIP-Seq(GSE58341)/Homer | LTA |
| ZSCAN22(Zf)/HEK293-ZSCAN22.GFP-ChIP-Seq(GSE58341)/Homer | LTA |
| DUX4(Homeobox)/Myoblasts-DUX4.V5-ChIP-Seq(GSE75791)/Homer | R848 |
| Esrrb(NR)/mES-Esrrb-ChIP-Seq(GSE11431)/Homer | R848 |
| FOXM1(Forkhead)/MCF7-FOXM1-ChIP-Seq(GSE72977)/Homer | R848 |
| IRF4(IRF)/GM12878-IRF4-ChIP-Seq(GSE32465)/Homer | R848 |
| TEAD3(TEA)/HepG2-TEAD3-ChIP-Seq(Encode)/Homer | R848 |
| ZKSCAN1(Zf)/HepG2-ZKSCAN1-ChIP-Seq(Encode)/Homer | R848 |
| ZNF317(Zf)/HEK293-ZNF317.GFP-ChIP-Seq(GSE58341)/Homer | R848 |
| Phox2b(Homeobox)/CLBGA-PHOX2B-ChIP-Seq(GSE90683)/Homer | SA |
| PSE(SNAPc)/K562-mStart-Seq/Homer | SA |
